# CellDepot: A unified repository for scRNA-seq data and visual exploration

**DOI:** 10.1101/2021.09.30.462602

**Authors:** Dongdong Lin, Yirui Chen, Soumya Negi, Derrick Cheng, Zhengyu Ouyang, David Sexton, Kejie Li, Baohong Zhang

## Abstract

CellDepot serves as an integrated web application to assist users in exploring single-cell RNA-seq (scRNA-seq) datasets and comparing the datasets among various studies through a user-friendly interface with advanced visualization and analytical tools. To begin with, it provides an efficient data management system that users can upload single cell datasets and query the database by multiple attributes such as species and cell types. In addition, the advanced query function incorporated in MySQL database system and its conditional filtering, allows users to quickly query and compare the expression of gene(s) across the datasets of interest. Moreover, by embedding the cellxgene VIP tool, CellDepot enables fast exploration of individual dataset in the manner of interactivity and scalability to gain more refined insights such as cell composition, gene expression profiles, and differentially expressed genes among cell types. In summary, the web portal allows large scale single cell data sharing, analysis and visualization for supporting decision-making, and encouraging scientists to contribute to the single-cell community in a tractable and collaborative way. Finally, CellDepot is released as open-source software to motivate crowd contribution, broad adoption, and local deployment for private data.

## Introduction

Since the first print of using next-generation sequencing technology to analyze the single-cell transcriptome in 2009 [1], this technology has massively ignited the interest in obtaining high-resolution characterizations of each cell’s transcriptome. Afterwards, an exponentially growing number of studies utilized this advanced scRNA-seq technology to investigate the expression changes among cells or groups from various sources such as cell lines, tissues, or species [2-5]. Meanwhile, substantial efforts have been invested in developing computational pipelines and tools to advance the analysis and visualization of large-scale scRNA-seq data [6-10]. With more scRNA-seq datasets have been published by community, it is urgent to launch some in-depth investigations of the associations of those identified biological variations with cellular heterogeneity. However, many datasets are from different studies or individual labs that are preprocessed in varied ways. Therefore, it is necessary to build a centralized space for managing, analyzing, and visualizing those datasets to yield comprehensive insights from this big data.

Several tools and databases have been built to store, analyze, and visualize scRNA-seq datasets, which allow scientists to compile and query the information at their hand. The ‘Human Cell Atlas’ consortium [11] leads an international collaboration to generates single-cell datasets from human body tissues, which are manually curated and processed by a uniform pipeline. ‘JingleBells’ provides single-cell data [12] focusing on immune cells. The ‘conquer’ database provides uniformly processed single-cell expression data to facilitate benchmarking of computational tools [13]. The ‘PanglaoDB’ database provides single-cell RNA-seq count matrices from public sequencing data in the National Center for Biotechnology Information Sequence Read Archive [14]. The ‘EMBL-EBI Single-Cell Expression Atlas’ provides uniformly processed data from submissions to ‘ArrayExpress’ [15]. The Broad Institute also offers a public ‘Single-Cell Portal’ (https://singlecell.broadinstitute.org/single_cell). Although those ‘atlas’ and portals have greatly facilitated the exploration of single-cell datasets from the database [16-20], they usually tackle certain aspects of managing, visualizing, or analyzing scRNA-seq datasets with limitations on both query and visualization functionalities. For comprehensive review of these portals, detailed feature comparison is outlined in supplementary Table S1.

We developed a database-backed single cell portal available at http://celldepot.bxgenomics.com with user-friendly interactive visualization and analysis capabilities, namely CellDepot to empower biologists and bioinformaticians to manage, explore, visualize, and compare scRNA-seq datasets in a comprehensive, flexible, and collaborative manner.

## Results and Discussion

### 1. Overview of CellDepot

CellDepot is a user-friendly centralized database platform that integrates the database management system along with the search engine. The web-based platform enables the share of scRNA-seq data and efficient communication among the community, which in turn encourages crowd contributions to the online scRNA-seq data portal. Currently, there are more than two hundred scRNA-seq datasets from eight species hosted on the portal for public access. Notably, CellDepot integrates with advanced single-cell transcriptomic data explorer to conduct all analytical tasks on the webserver while presenting interactive results on the webpage through leveraging modern web development techniques (Figure 1a).

**Figure 1.**
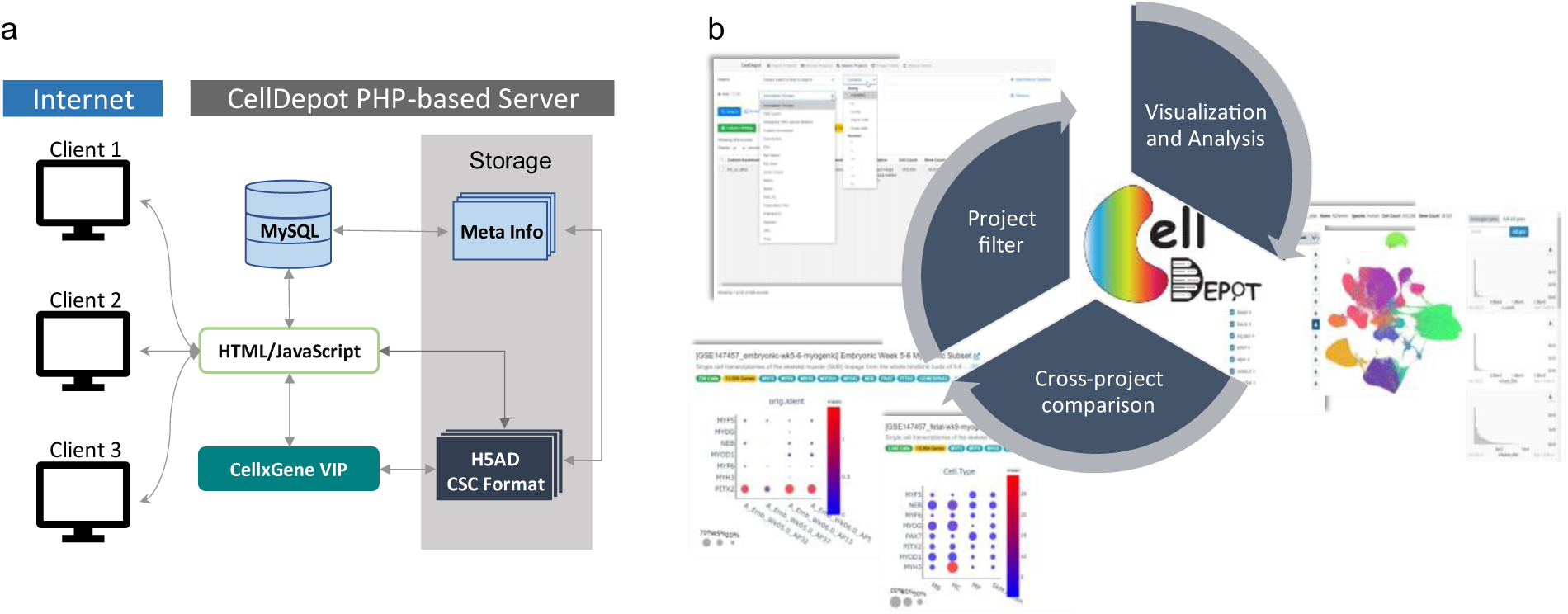
CellDepot portal overview. (a) The architecture of CellDepot. (b) Functional structure of CellDepot.

Highly unified space for managing, exploring, and analyzing comprehensive single-cell datasets is one of the significant strengths of CellDepot. Nowadays, while a few web-based portals for both visualizing and analyzing scRNA-seq datasets are available, they have limitations on scalability and capability in comparing among various datasets to meet increasing demands on examining single-cell RNA-seq datasets. CellDepot employs the MySQL relational database management system, a robust scRNA-seq data visualization tool cellxgene [21], and its versatile plugin cellxgene VIP [22] to allow project selection and filtering, scRNA-seq data visualization and analysis, and cross-project comparison of targeted genes as illustrated Figure 1b.

### 2. Features of CellDepot

There are mainly three functions built in the platform: project search/filter, visualization and analysis, and cross-project comparison, which will be discussed thoroughly in the following sections.

#### a. Data filtering

scRNA-seq datasets are stored as h5ad files in the compressed sparse column (CSC) format and managed efficiently in the CellDepot together with the metadata table of projects. For each dataset, some primary metadata fields are inputted by users and can be updated as needed. The main CellDepot application consists of five functionalities:

- **Importing projects** provides a user-friendly way to upload processed public or private datasets in h5ad format.
- **Browsing projects** displays a quick overview of all datasets in CellDepot with a quick search function and customized column setting such that users can easily find the projects of interest, personalize the look-n-feel of the project table and export the customized project spreadsheet.
- **Searching projects** enables advanced query of projects by joining multiple search conditions with the logical operators.
- **Project filters** refines the matched datasets by simply selecting ‘Year’ and/or ‘Species’. It is a user-friendly feature for a first-time user who is not familiar with the content of the database.
- **Searching genes** allows users to input any gene(s) of interest and generate the cross-dataset plots for those targeted genes.

Based on available meta information stored in the h5ad files, multiple dataset attributes are extracted for searching and filtering, such as species, cell type annotation, published year, etc. As shown in Figure S1, there are six datasets when searching by ‘Species is Human’ and ‘Annotation Group contains Neuron’.

It is desirable to include all possible abbreviations and synonyms of genes or cell types in the database to make search more flexible. In reality, CellDepot has not enumerated and standardized all possible terms used to name cell types in the query function beyond metadata collected from h5ad files. In the future, a comprehensive mapping table of terms based on cell ontology could be implemented to standardize keywords for searching. Alternatively, user can use advanced search function to query a cell type by multiple synonyms, e.g., searching by ‘Annotation Groups contains OPC’ or ‘Annotation Groups contains oligodendrocyte’ to identify datasets containing oligodendrocyte cell type.

#### b. Data visualization and analysis

Users can launch a cellxgene VIP instance (such as dataset GSE140231) [23] as shown in Figure 2 to explore a dataset interactively. Cellxgene VIP is based on the cellxgene platform where users can quickly overview meta information and cell embedding after dimensional reduction (e.g., TSNE or UMAP). In addition, users can use lasso selection from embedded cell maps to interactively select any cell cluster(s) as a group for refined analysis via a rich set of visualization and analysis functions provided in cellxgene VIP.

**Figure 2.**
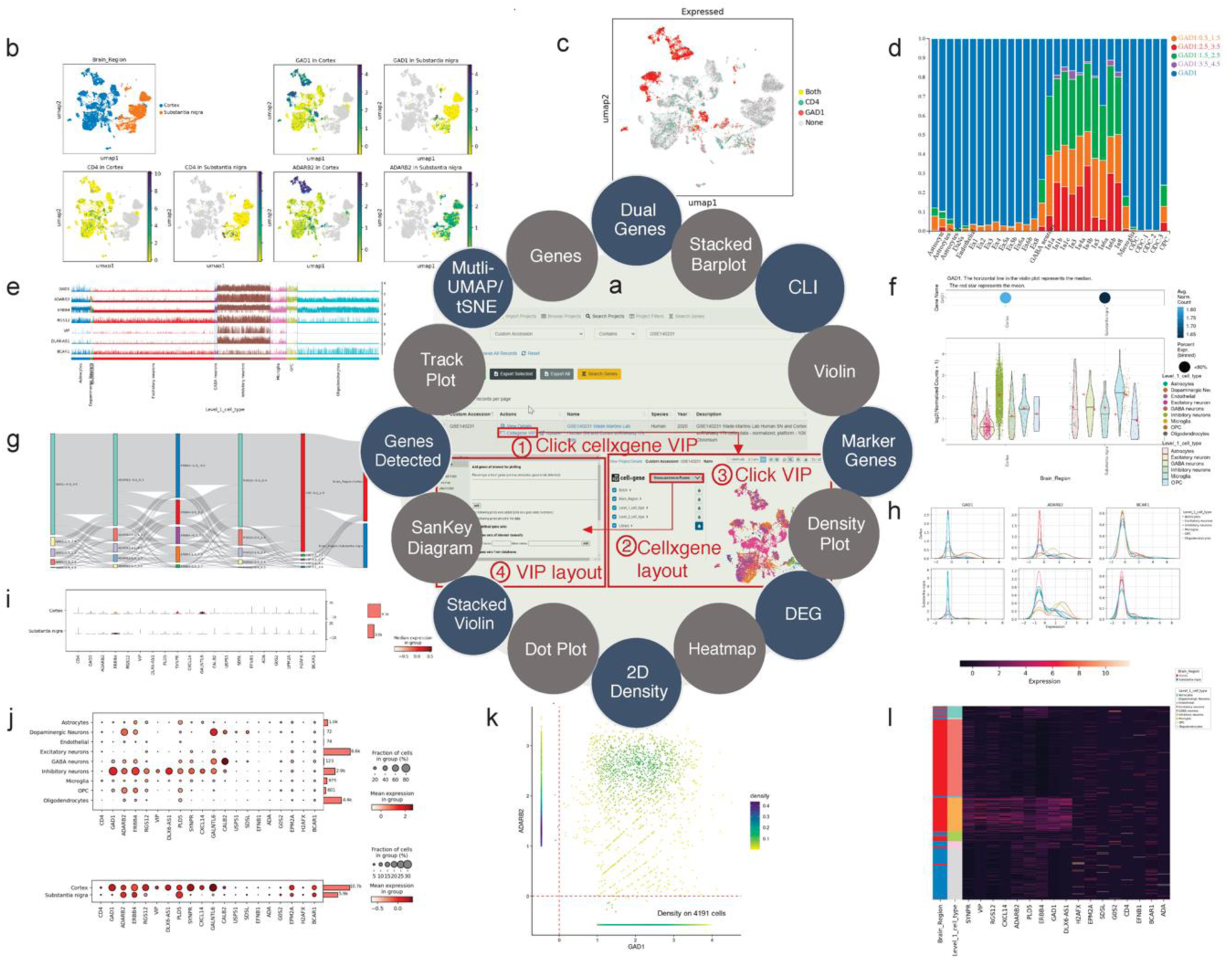
Exploration of visualization and analysis functions in cellxgene VIP on dataset GSE140231 [23]. (a) **Cellxgene VIP**, providing a plugin ecosystem of interactive data visualization, can be launched within four steps after querying ‘custom accession contains GSE140231’. (b) **Mutli-tSNE/UMAP plot** shows the differential expression of genes on split brain regions. (c) **Dual-gene plot** demonstrates the expression of CD4 and GAD1 on the selected embedding layout. (d) **Stacked barplot** indicates the fraction of cell distribution of the selected gene over different cell types. (e) **Trackplot** represents the distribution of gene expression across individual cells in annotated clusters. (f) **Sub-grouped violin plot** shows the GAD1 gene expression across the groups of cell types and subgroups of the brain regions. (g) **Sankey diagram** illustrates the inter-dependent relationship of annotated clusters based on the selected genes. (h) **Density plots** show expression of marker gene across varied cell types and split across the brain regions. (i) **Stacked violin plot** highlights the selected cell markers over cell types. (j) **Dot plots** highlights the gene markers over different annotated clusters. (k) **2D-density plot** illustrates the expression relationship of two genes (GAD1 and ADARB2). (l) **Heatmap** of selected marker genes in various cell types.

For example, user can go deep-dive to explore and visualize the expression of gene(s) across the cluster of cells under various conditions (Figure S2). As shown in Figure S2a, two cell groups from Astrocytes (1036 cells) and Oligodendrocytes (4417 cells) are selected. By running differential analysis with one of the built-in statistical methods such as Welch’s t-test, we detected 1578 differential expressed genes (DEGs), including 715 up-regulated and 853 down-regulated genes in astrocytes compared to oligodendrocytes (Figure S2a). The expression of the top four DEGs among the cell types indicates that gene MBP, ST18 and RNF220 are expressed explicitly in oligodendrocytes, while gene PITPNC3 is expressed mainly in astrocytes and endothelial cells (Figure S2b). In the future, we plan to add other multi-omics data modalities, which can be incorporated and integrated with scRNA-seq, such as spatial transcriptomics and scATAC-seq data.

#### c. Cross-dataset query

Besides the function to visualize and explore individual datasets, CellDepot also allows users to query and compare the expression of genes of interest across multiple datasets to understand their cellular heterogeneity. As shown in Figure 3, users can explore the expression of skeletal muscle marker genes PAX3, PAX7, PITX2, MYF5, MYF6, MYOD1, MYOG, NEB, and MYH3 during human myogenic cells of development and differentiation from hx-protocol 4-6 weeks to fetal 9-18 weeks from datasets whose custom accession starting with GSE147457 [24]. Gene expression level in SkM cells highlighted by grey boxes in of Figure 3e decreases around 40% from week 4 to week 5 with hx-protocol until level-off at week 6. The expression level of genes in fetal stage over week 9 also shows the same trend, especially the gene expression level drops dramatically starting from week 12.

**Figure 3.**
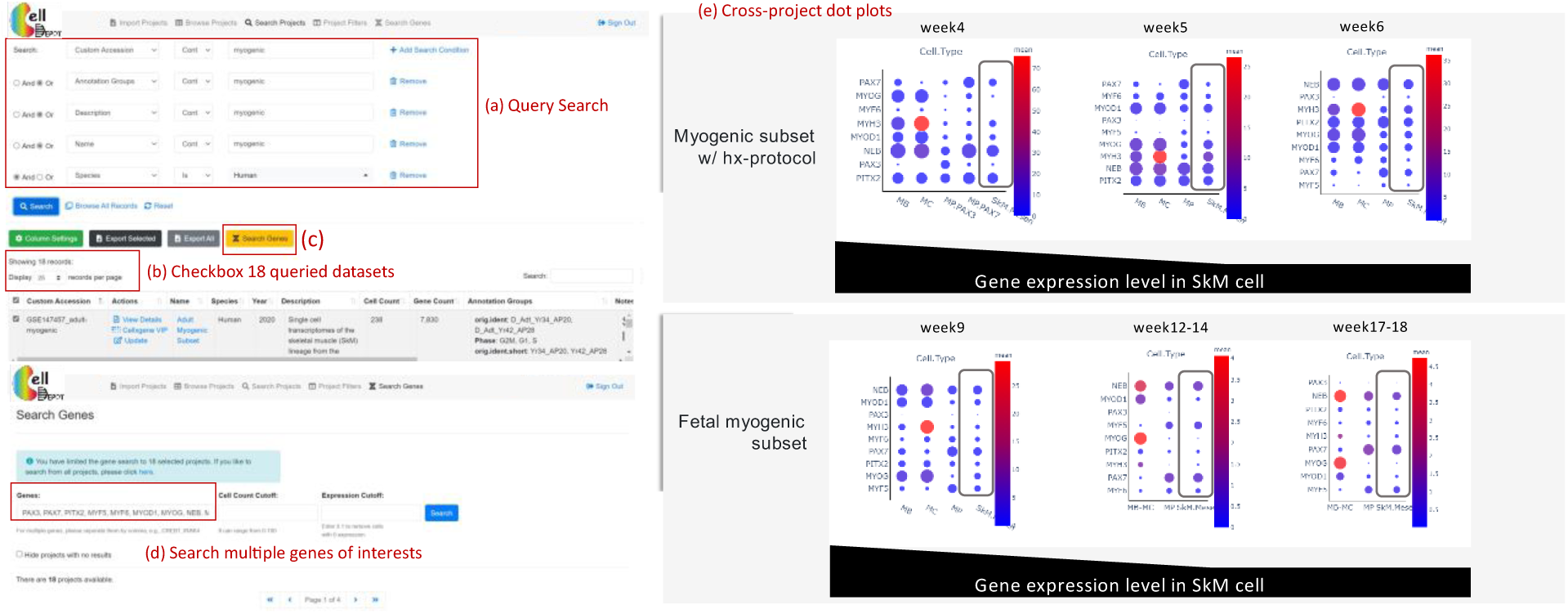
Cross-project view of the expression of genes among skeletal muscle development and differentiation times. (a) Search and (b) select the targeted datasets whose custom accession starting with GSE147457 [24]. Navigate to (c) ‘Search Genes’ page and (d) search genes of interest. (e) The summary of interactive plots from CellDepot that shows the cross-project comparison of gene expression level under varied conditions.

In summary, CellDepot is an easy-to-use and elaborative web portal for the exploration of the scRNA-seq datasets and data analysis results such that it allows biologists to easily access and reuse the rapid-increased scRNA-seq datasets in a highly scalable and interactive manner.

## Materials and Methods

### Sources of annotation and metadata

The original metadata information of each single cell RNA-seq dataset is retrieved from h5ad file, which is a preferred way of sharing and storing an on-disk representation of anndata object [6]. When importing the dataset to the system, user inputs additional metadata information as shown in Figure S5. Both metadata are collected and stored in a MySQL database table that is presented at http://celldepot.bxgenomics.com.

### Data format, availability and preparation

CellDepot requires scRNA-seq data in h5ad file where the expression matrix is stored in CSC (compressed sparse column) instead of CSR (compressed sparse row) format to improve the speed of data retrieving. For example, designating genes as columns in the h5ad file creates the interactive plot five times faster than as rows. Just in case, we provide sample scripts to help users generate h5ad files. Having gene expression matrix, metadata, and layout files, users can easily combine and convert their data to h5ad file by following this R script on https://github.com/interactivereport/CellDepot/blob/main/toH5ad.R. In the case of lacking layout file, users can also create h5ad file by following the Jupyter notebook https://github.com/interactivereport/CellDepot/blob/main/raw2h5ad.ipynb with custom python script tailored to their own data. Categorical features extracted from a h5ad file are shown in the ‘;annotation groups’; column of the table on CellDepot home page, while the numerical features are shown as the distributions in the rightmost panel on cellxgene VIP (Figure S3).

### CellDepot platform and installation

The public version of CellDepot web portal is hosted at the web site, http://celldepot.bxgenomics.com and is implemented with MySQL database, an advanced search engine, and powerful interactive visualizing tools that allow users to explore attributes of datasets as well as scRNA-seq analysis results. Also, users can intentionally select single-cell RNA-seq datasets on the web interface by simply browsing the online dataset table or applying advanced search to perform the cross-dataset comparison. Moreover, CellDepot also provides comprehensive data analysis tools via an embedded interactive visualization plugin. To host private datasets, local instance of CellDepot on Unix server can be installed by following the guide here, https://celldepot.bxgenomics.com/celldepot_manual/install_environment.php.

### Data import on user’s server

The prepared h5ad files are required to copy to a folder defined in the configuration file, e.g., /data/celldepot/all_h5ad_files/. Afterwards, users can navigate to the CellDepot home page, click ‘Import Project’ at the top menu, then ‘Download Example ile’ to fill in meta information of datasets into the downloaded template for submission. After the metadata file is uploaded, CellDepot will automatically convert the dataset to CSC format if needed through a cron job (Supplementary Tutorial Section 5). To explore the detail of imported datasets, users can enter ‘Browse Projects’ page and then search these datasets by user assigned accessions in the metadata file.

### CellDepot API (Application Programming Interface)

The CellDepot API web service provides a direct way to generate figures for users to share or embed in web page. For example, the following URL will generate a gene expression violin plot across cell clusters for IRAK4 gene for the data set with ID equaling one, https://celldepot.bxgenomics.com/celldepot/app/core/api_gene_plot.php?ID=1&Genes=IRAK4&Plot_Type=violin&Subsampling=0&n=0&g=0&Project_Group=CLUSTER. The complete format of the URL and explanation of parameters are detailed in the web page, https://celldepot.bxgenomics.com/celldepot_manual/api_gene_plot.php.

## Code availability

The source code, local installation guide and complete tutorial of visualization and analysis tool are provided at https://github.com/interactivereport/CellDepot. With broad adoption and contribution in mind, CellDepot is released under the MIT License.

## Author Contributions

Dongdong Lin: Conceptualization, Methodology, Manuscript – writing & review

Yirui Chen: Testing, Manuscript – writing, review & editing

Kejie Li: Conceptualization, Testing, Manuscript – review & editing

Soumya Negi: Methodology, Testing, Manuscript – review & editing

Derrick Cheng: Web application development, Manuscript – review

Zhengyu Ouyang: Data curation, Manuscript - review

David Sexton: Conceptualization, Manuscript - review

Baohong Zhang: Conceptualization, Testing, Methodology, Manuscript – writing, review & editing

## Acknowledgements

The authors are grateful and indebted to BioInfoRx, Inc. for the management of server and storage to host the public web site.

## Supplementary Information

### Supplementary Tables

**Table S1.**
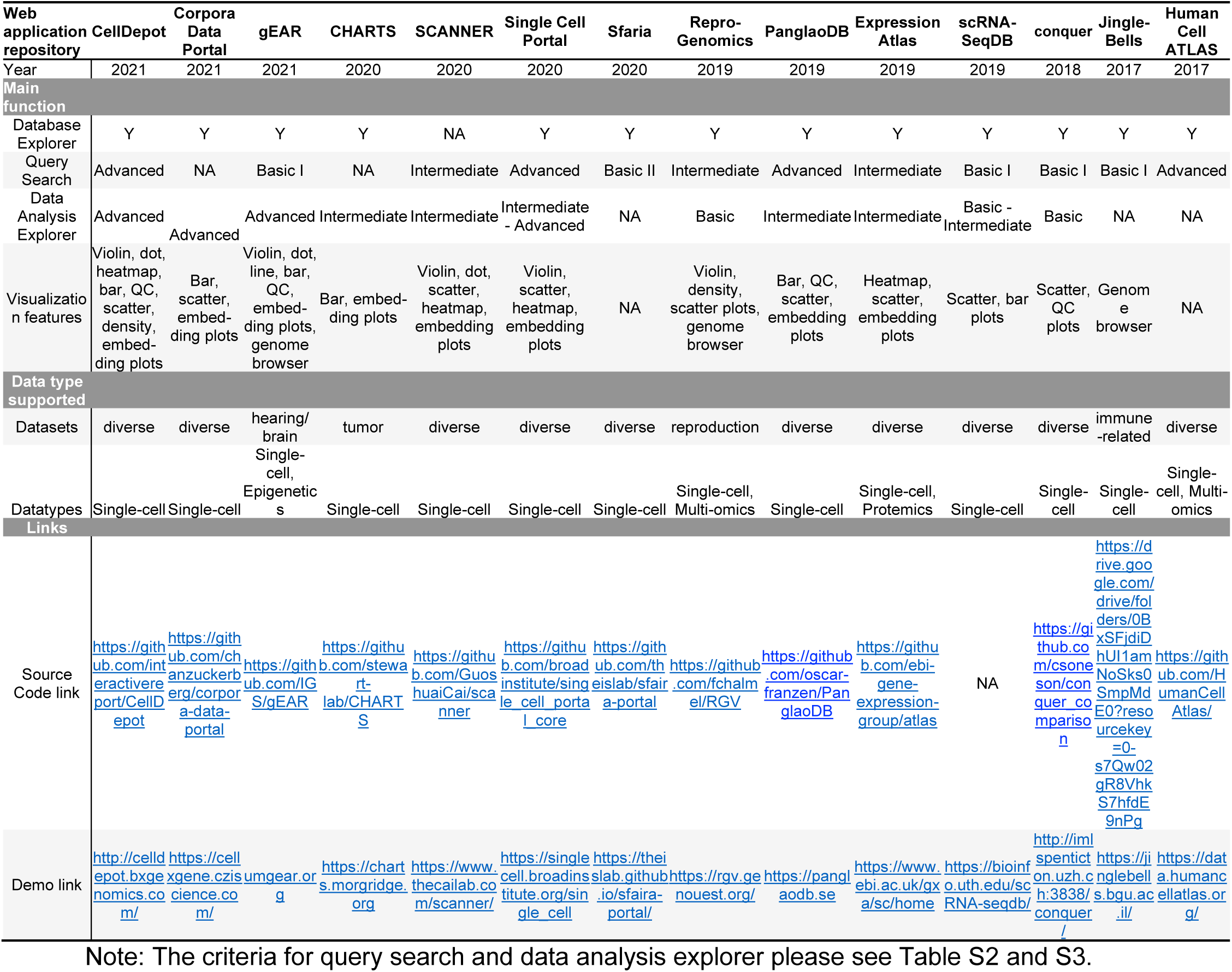
Comparison matrix of web portal tools

**Table S2.** Criterion for query search

| Query Search | Keyword Search | Multiple Object Search | Category Filters |
| --- | --- | --- | --- |
| Basic | Y |  |  |
| Intermediate | Y |  | Y |
| Advanced | Y | Y | Y |

**Table S3.** Criterion for data analysis explorer

| Data Analysis Explorer | Visualization and Analyze<br>scRNA-seq Data | Visualization and Analyze<br>scRNA-seq Data<br>Analyze Gene Expression | Customize Displays |
| --- | --- | --- | --- |
| Basic | Y |  |  |
| Intermediate | Y | Y |  |
| Advanced | Y | Y | Y |

**Table S4.**
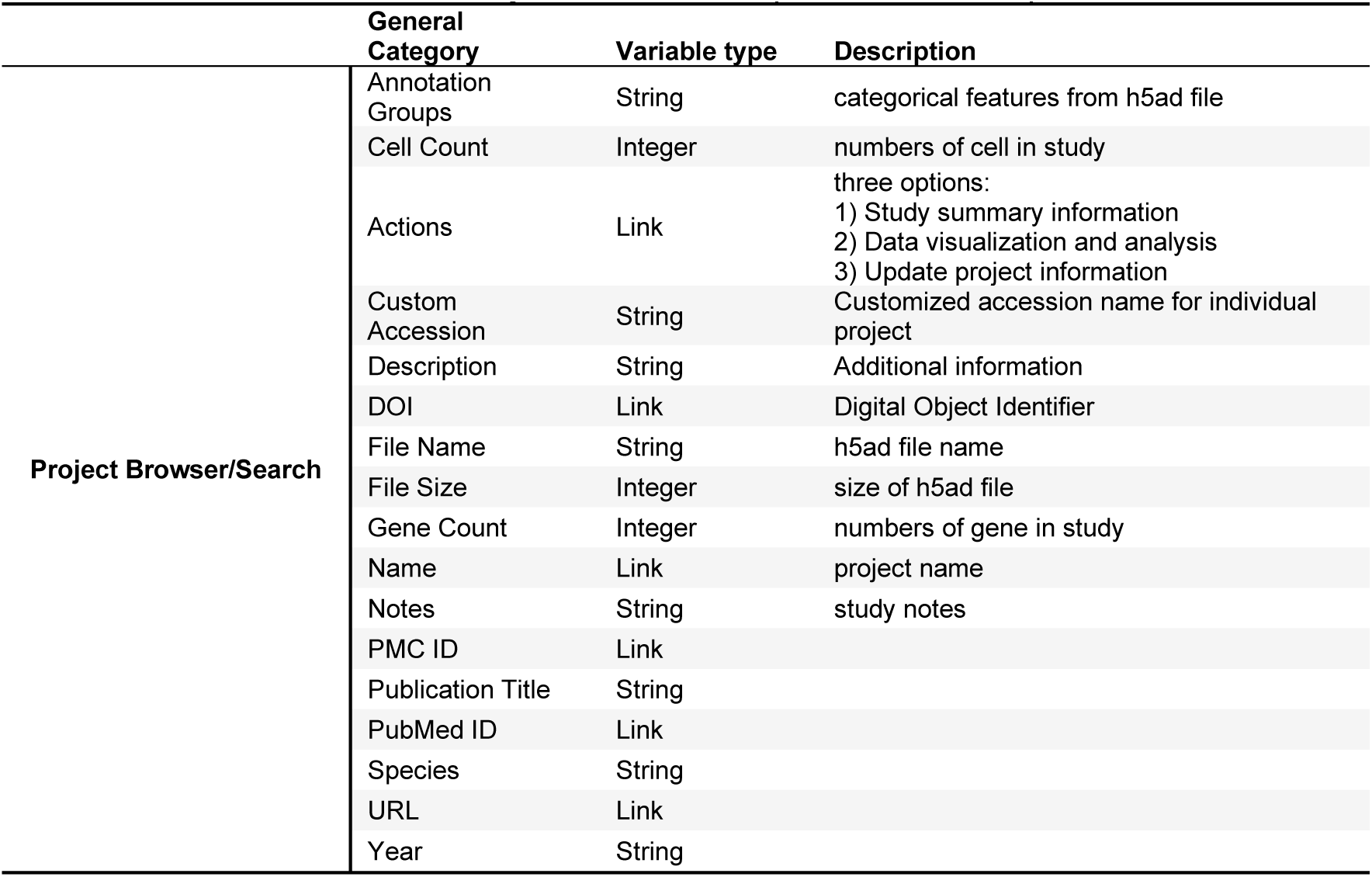
Project metadata captured in CellDepot

### Supplementary Figures

**Figure S1.**
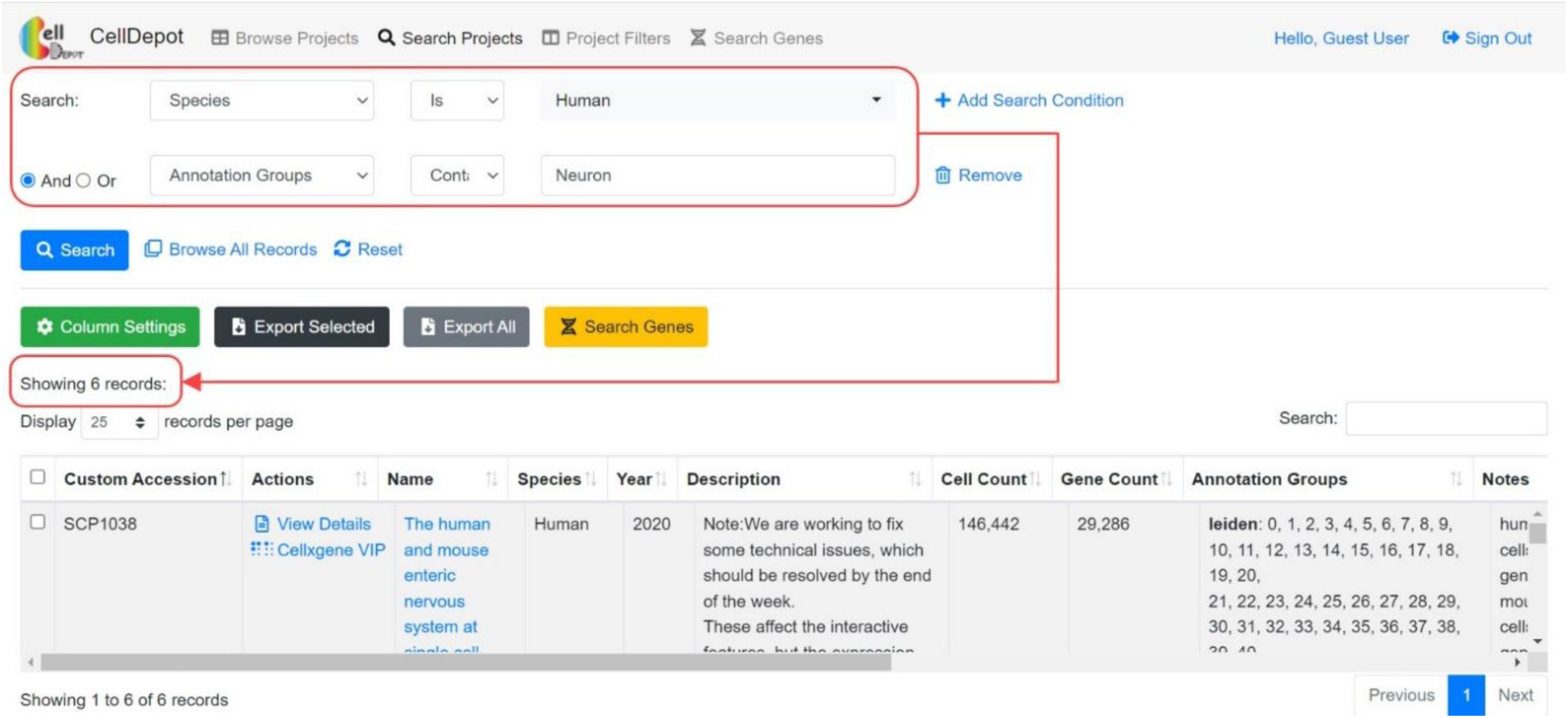
Data Filtering. Query search of ‘Species is Human’ and ‘Annotation Groups contains Neuron’ brings about nine datasets of interest.

**Figure S2.**
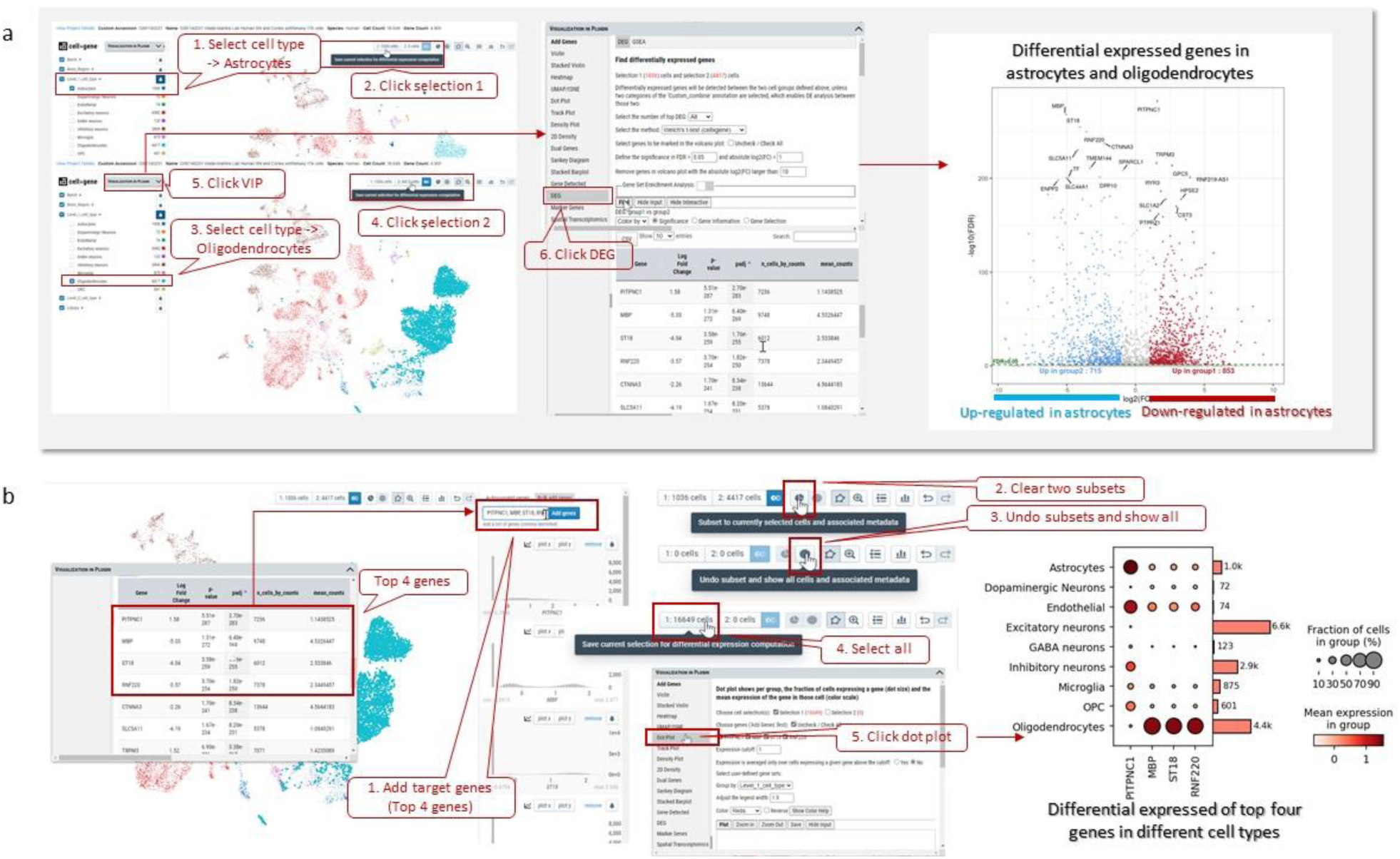
Exploration of differential expressed genes in dataset GSE140231 through cellxgene VIP [1]. (a) Differential expressed genes in astrocytes and oligodendrocytes. (b) The expression of top four genes in different cell types.

**Figure S3.**
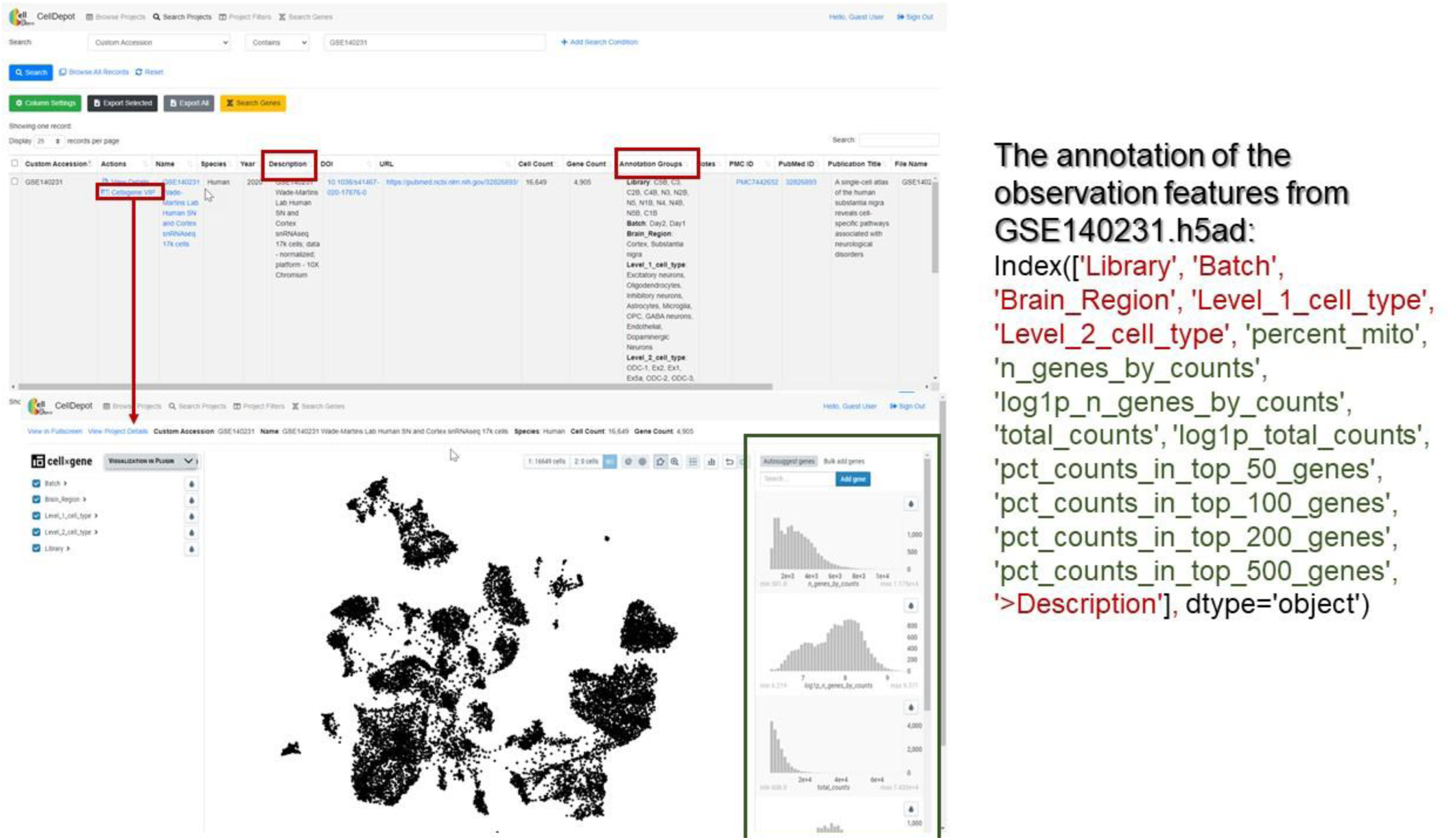
The exploration of observation features of dataset GSE140231. Red-marked categorical features are shown on CellDepot Project page (highlighted by red framed box), the numerical features marked in green color can be visualized as distribution plots on the rightmost panel in cellxgene VIP tool, which is highlighted by green box.

### Supplementary Tutorial

#### 1. Introduction

CellDepot is database management system integrated with management system, query searching and data visualization tools [2, 3] for scRNA-seq datasets, which can be accessed by the link http://celldepot.bxgenomics.com. This is a supplemental tutorial provides a detailed guide.

**Figure S4.**
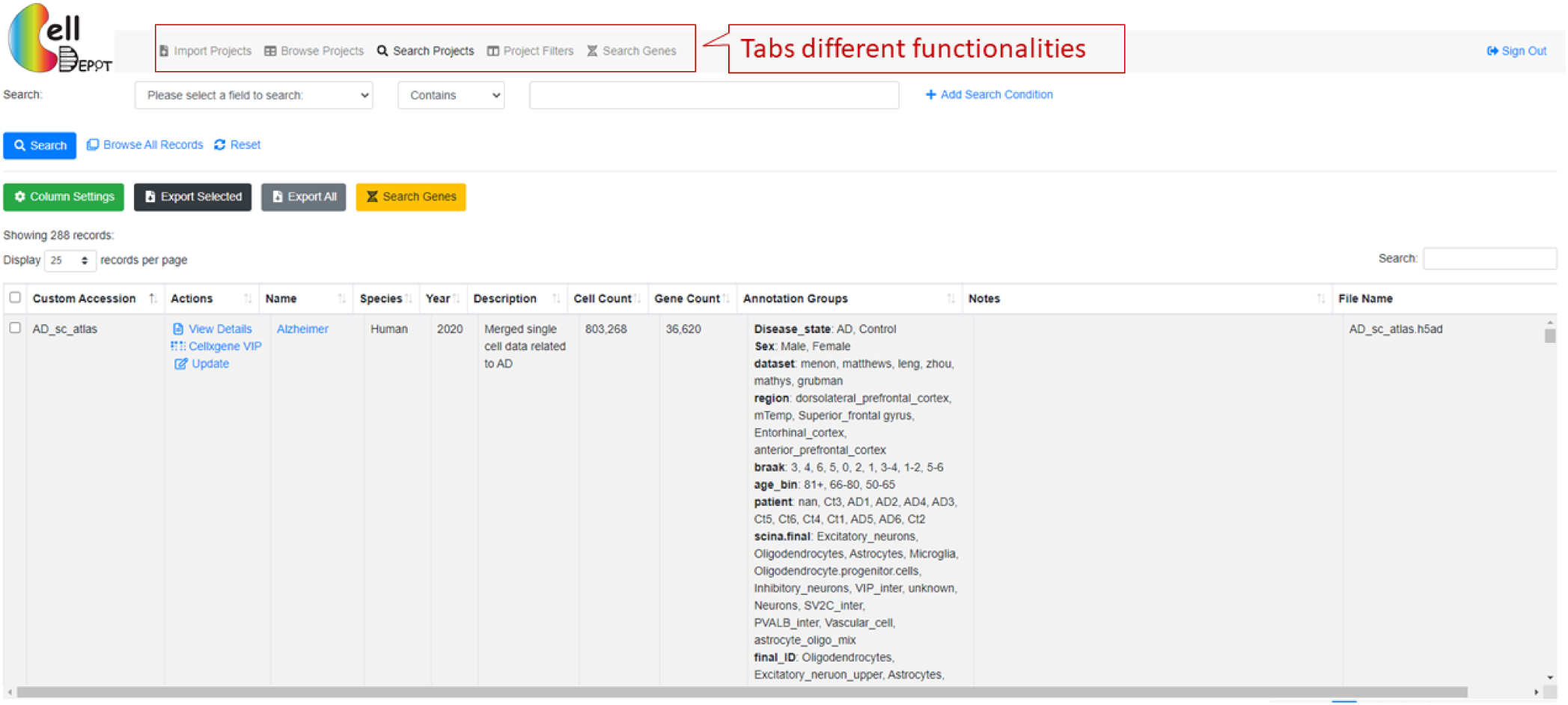
CellDepot website.

The interface contains multiple tabs, corresponding to import and/or select objects in CellDepot scRNA-seq database, that can be accessed on top panel of the webpage. Users can upload their own dataset or explore the existing datasets for visualization and analysis.

#### 2. Upload Projects

To upload new projects in CellDepot database, two files are required: 1) .h5ad files and 2) project information in csv format. Detailed formatting guidance can be found by ‘Download Example ile’ hyperlink on webpage. In addition, two cellxgene VIP launch methods are provided: standard and preload in memory. Standard mode is for the first-time imported datasets, while preload in memory should be chosen when users update the meta information of datasets.

After the projects are submitted, CellDepot will automatically analyze the datasets. To explore the detail of uploaded datasets, users can navigate to ‘browse projects’ page and then search the imported datasets by the customized accession number.

**Figure S5.**
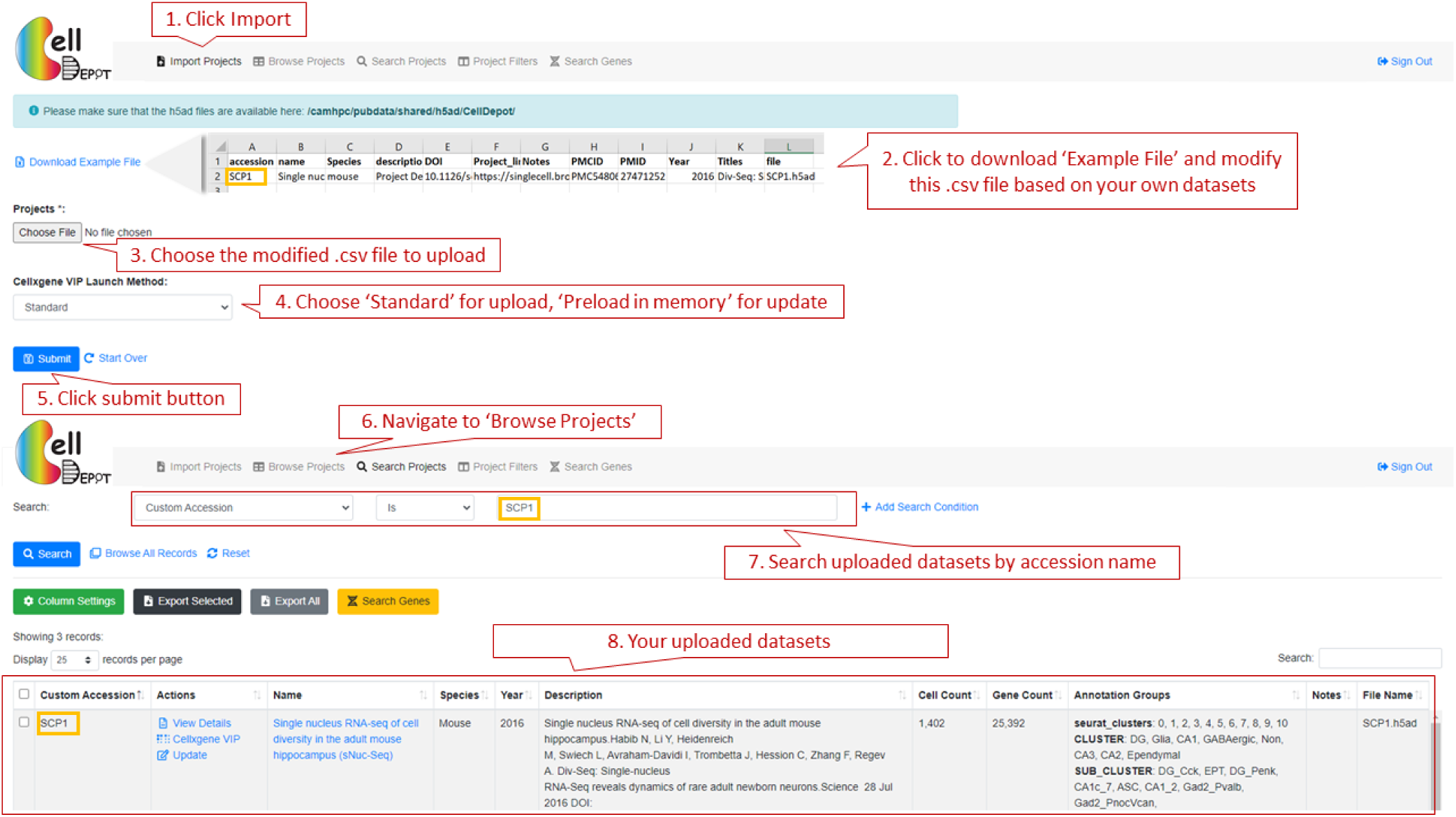
Workflow of how to import personal datasets.

#### 3. Browse Projects

##### 1. Search Projects

This function allows the user to search any targeted interests, which also can be accessed through search projects on the top panel of the webpage. Users is allowed to select the projects by 17 attributes: annotation groups, cell count, cellxgene VIP launch method, Custom accession, description, DOI, file name, file size, gene count, name, notes, PMC ID, Publication Title, PubMed ID, Species, URL, Year. These 17 fields can also be (partially) displayed on the webpage through ‘column setting’ on the webpage. Users can also search projects by the keywords via the search function on the right of the webpage. In addition, by ‘column setting’, users can set up the customized layout of targeted projects; thereby exporting to csv format.

For each project, users can view the datasets information, visualize data analysis, and conduct update through clicking on “View Details”, “cellxgene VIP”, and “Update” links, respectively.

**Figure S6.**
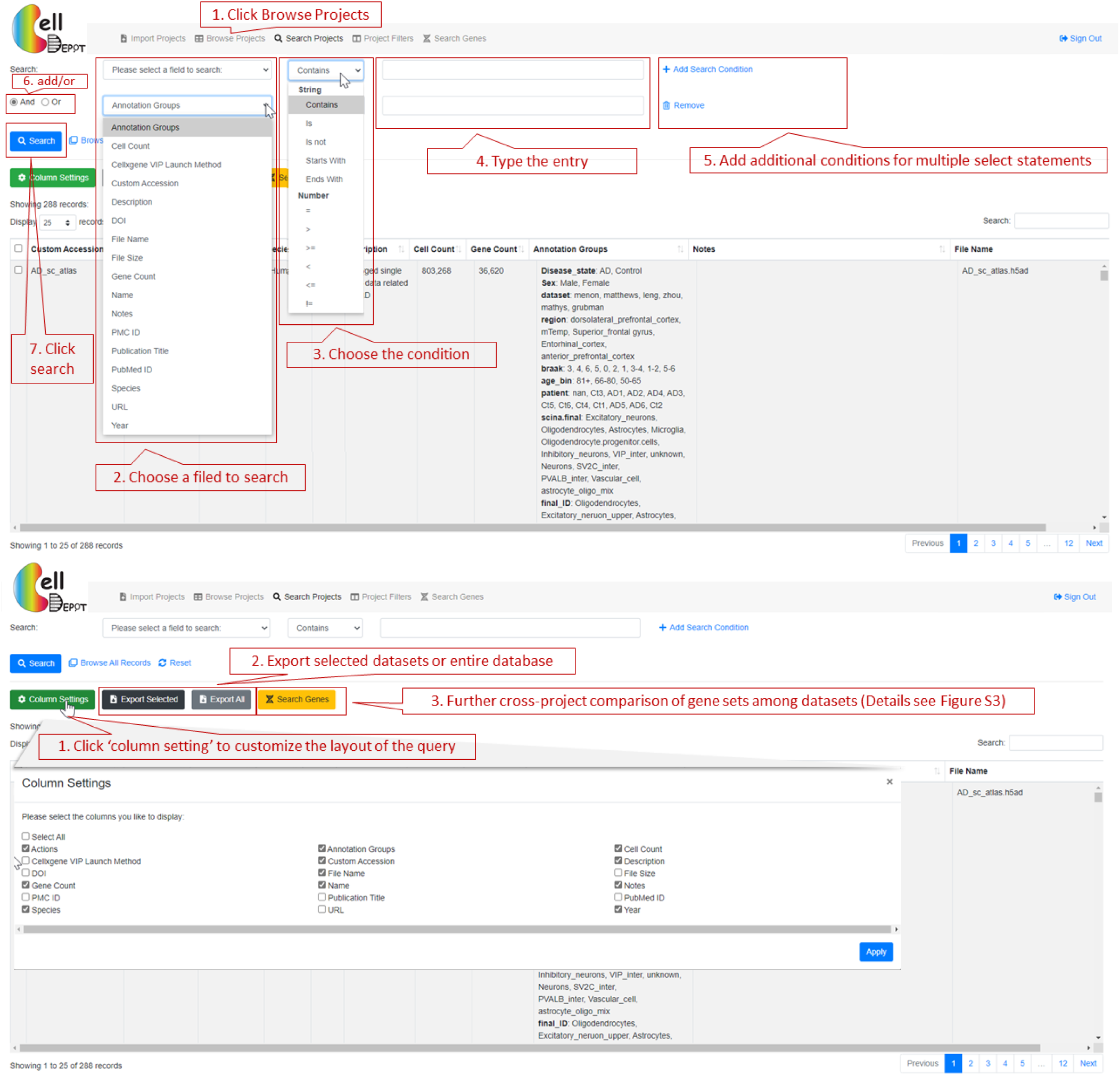
Workflow of how to search query on ‘Browse Projects’ page.

##### 2. Case Study 1

Cross-project comparison of skeletal muscle marker genes PAX3, PAX7, PITX2, MYF5, MYF6, MYOD1, MYOG, NEB, and MYH3 among the datasets whose species is human and cell type is myogenic.

**Figure S7.**
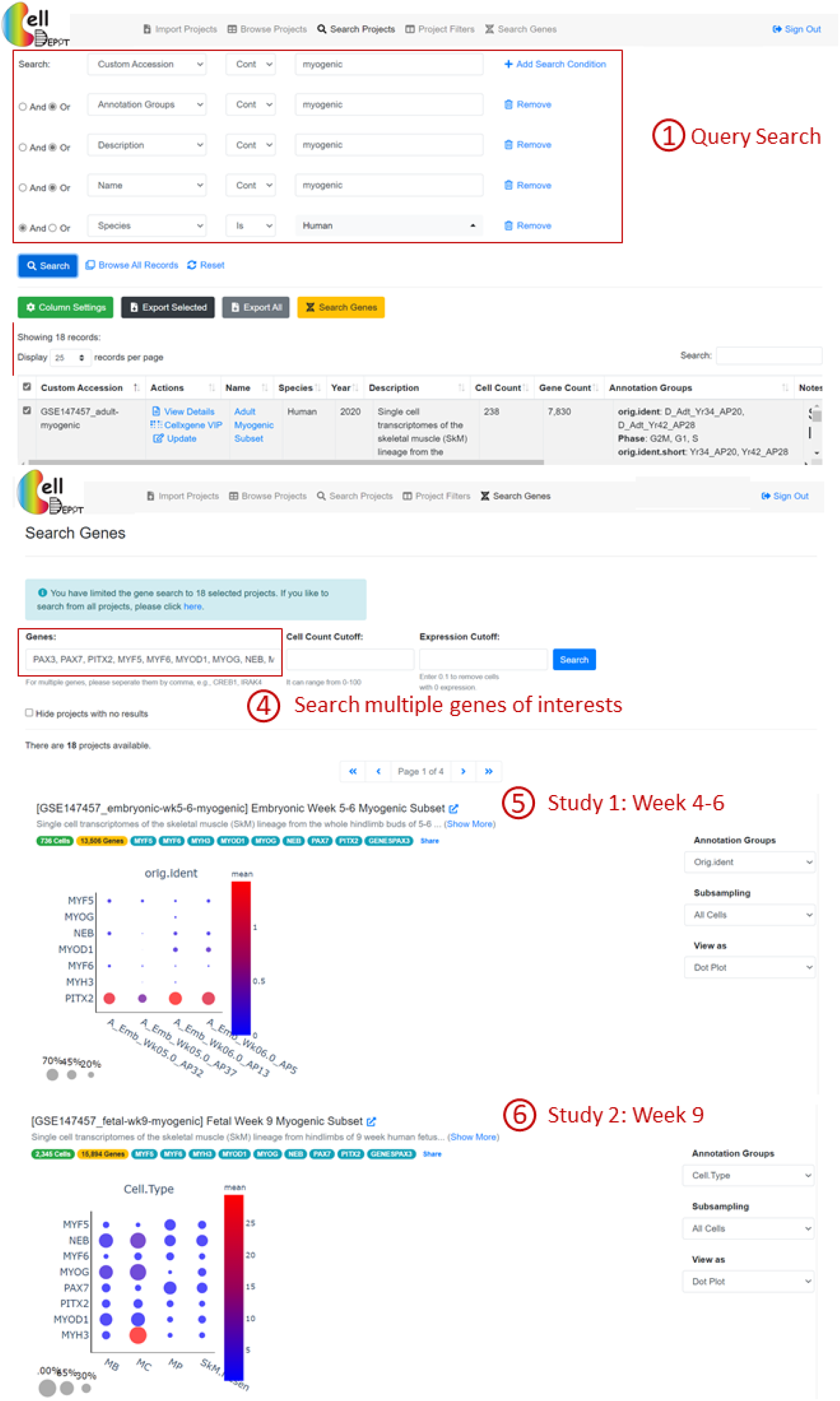
Workflow of how to conduct the cross-project comparison of gene sets among the selected datasets.

##### 3. Project Filters

This page provides the matched datasets by simply clicking the categories. It is a first-time user-friendly functionality as users may not be familiar with the content of the database. The advance search function is the same as that on the ‘Browse Projects’ page (Details see igure S4).

**Figure S8.**
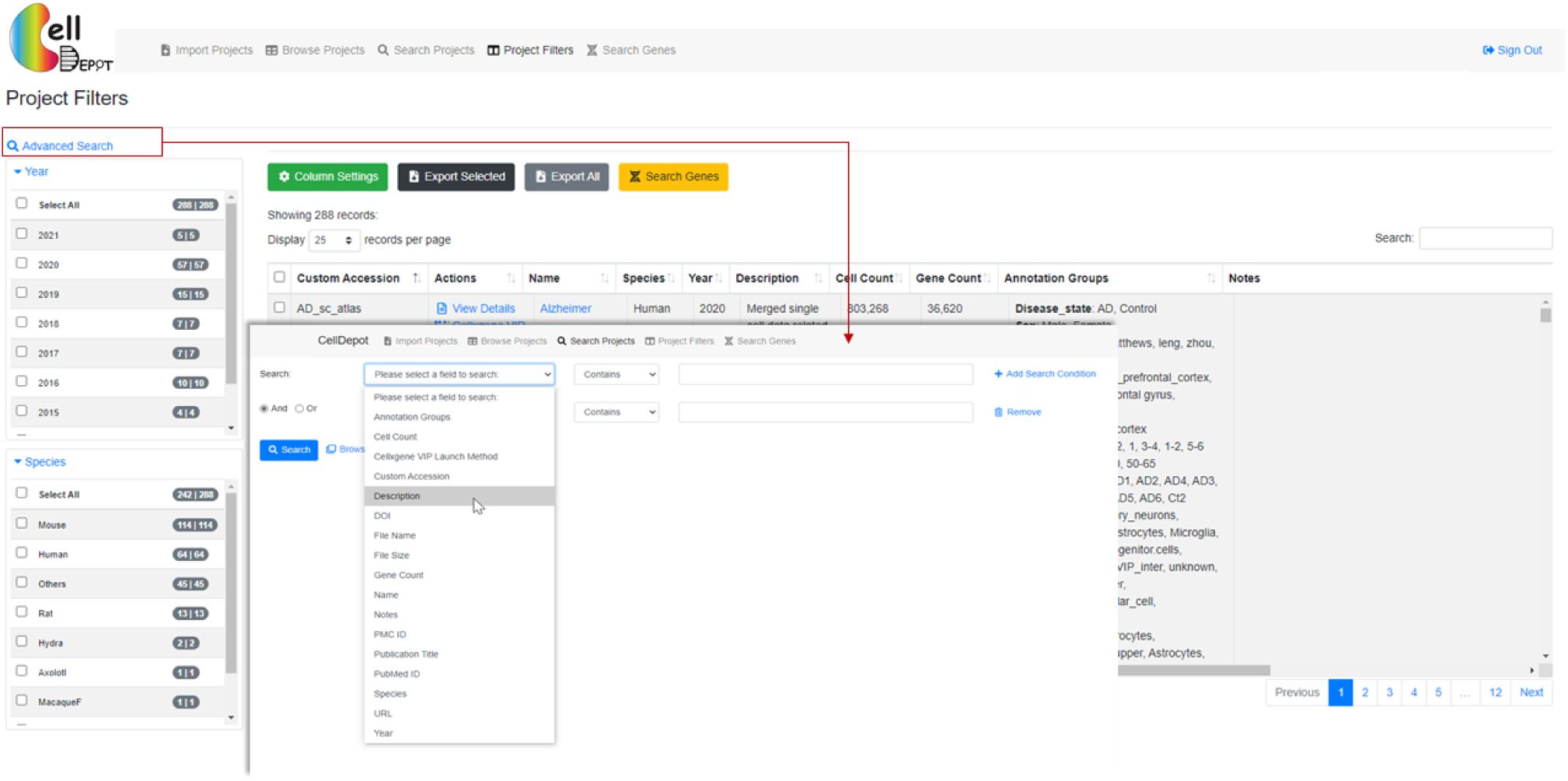
The layout of ‘Project Filters’ page.

#### 4. Visualize Datasets

##### 4.1 View Details

The datasets information consists of project summary and annotation groups. The project summary is provided by each user when uploading projects. The information of annotation groups is retrieved from uploaded .h5ad file.

**Figure S9.**
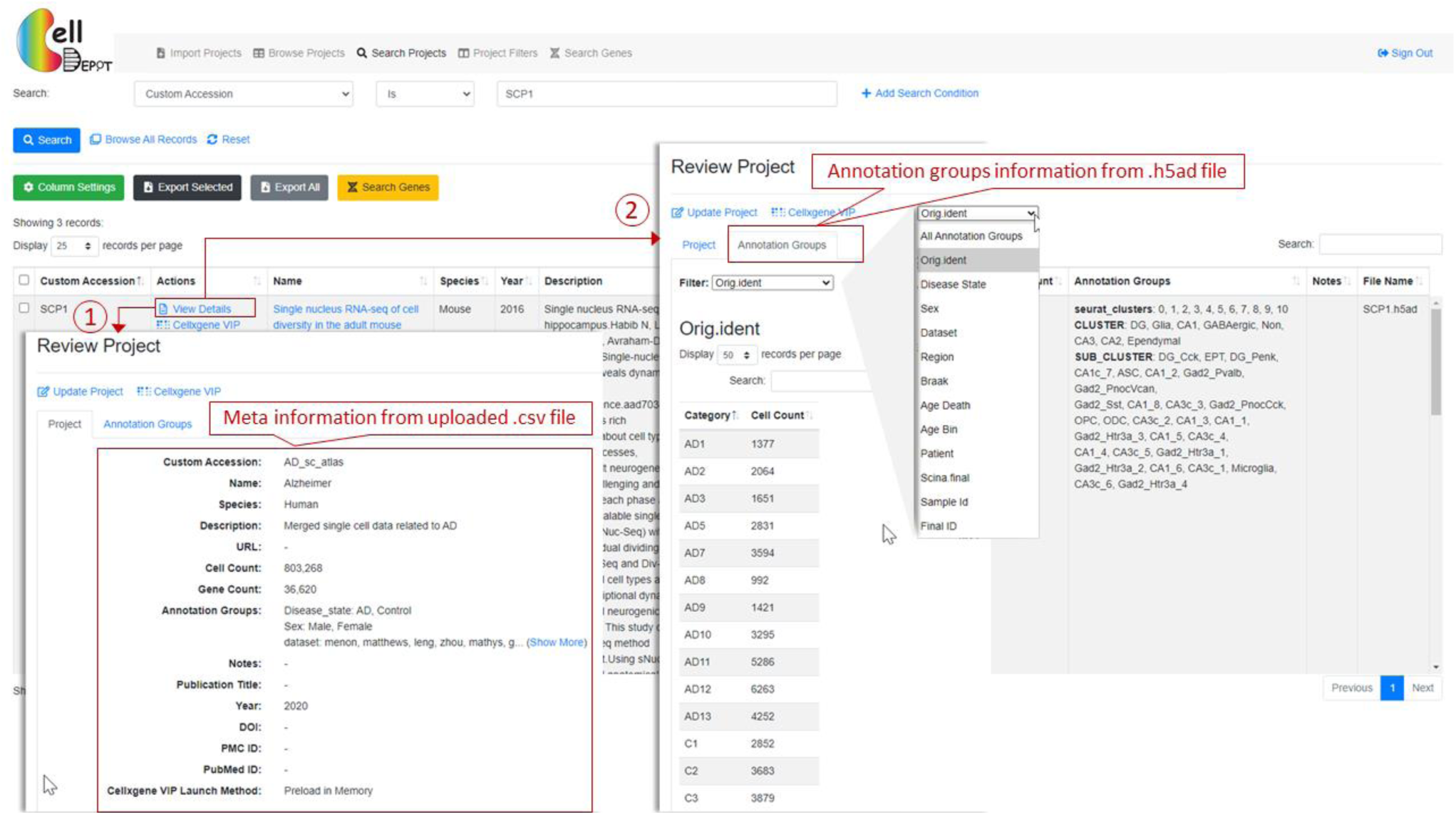
Visualization of details of datasets.

##### 4.2 Update

Project summary information can be updated on ‘Browse Project’ page with ‘Preload in Memory’ cellxgene VIP launch method via click ‘Update’ hyperlink.

**Figure S10.**
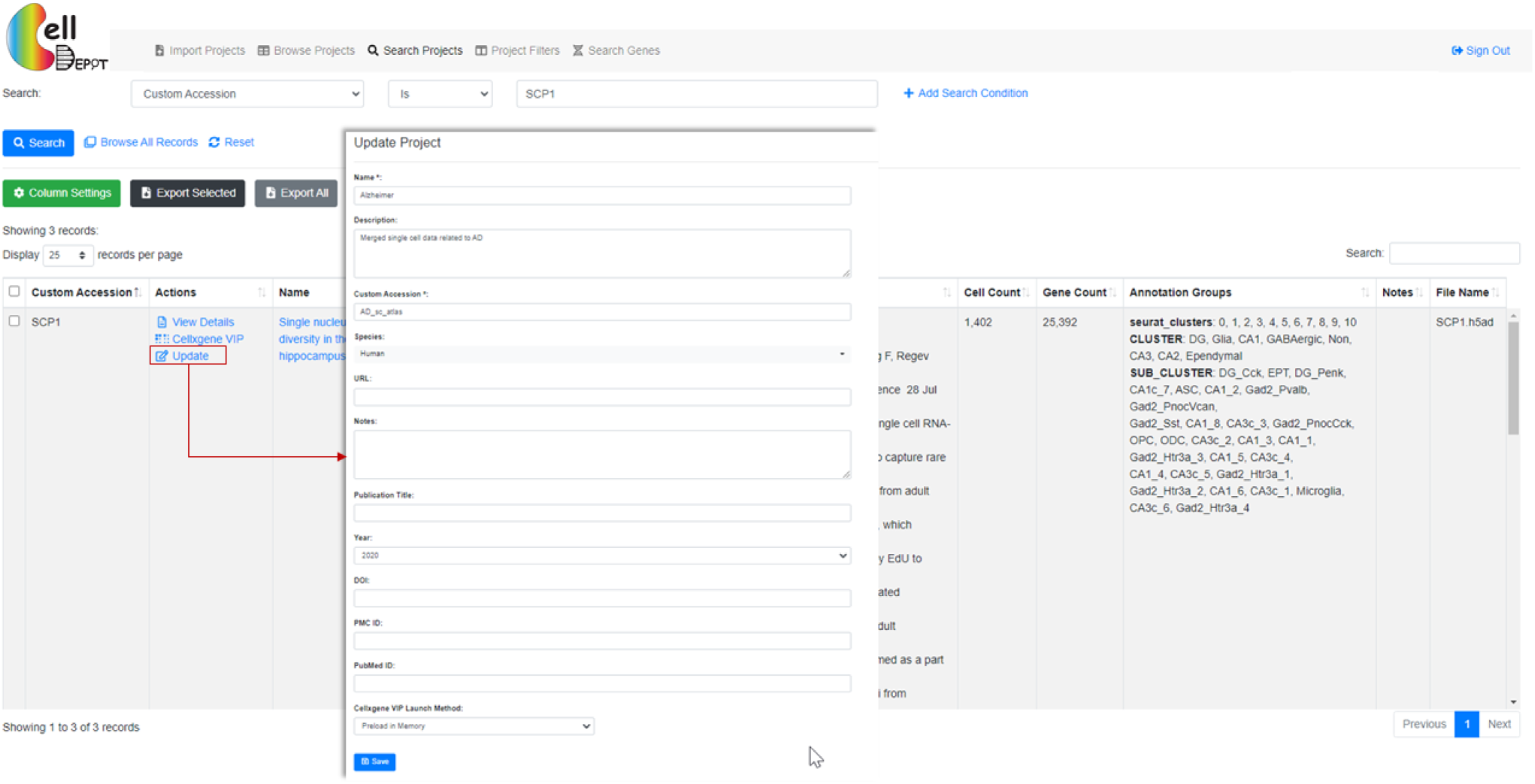
Update project on ‘Browse Project’ page.

##### 4.3 Data Visualization and Analysis

CellDepot is not only a database management system, but also a web portal to visualize the scRNA-seq dataset. Here, we embed cellxgene and cellxgene VIP in CellDepot. By clicking ‘Cellxgene VIP’, data analysis results can be visualized. Detailed guides of cellxgene and cellxgene VIP, please go to https://github.com/chanzuckerberg/cellxgene (cellxgene) and https://github.com/interactivereport/cellxgene_VIP (cellxgene_VIP).

**Figure S11.**
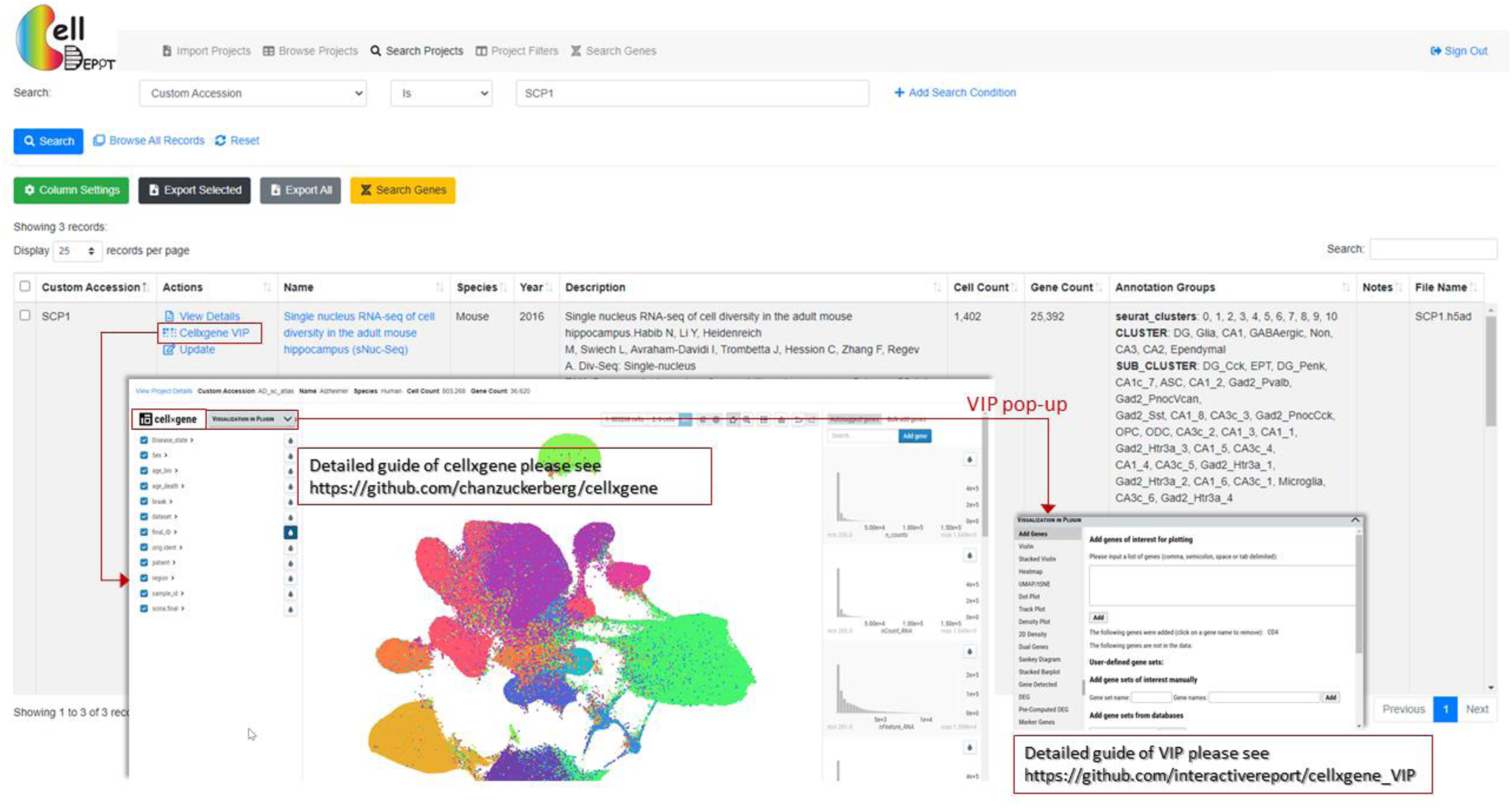
See visualization of selected datasets.

#### 5. Search Genes

This tab allows searching on targeted genes with cell count cutoff and expression cutoff. The search outcome provides users every project contains the targeted genes. Each project displays a link to project page and a figure plot if applicable. This plot can be either violin plot or dot plot shows the gene expression level in each annotation groups.

**Figure S12.**
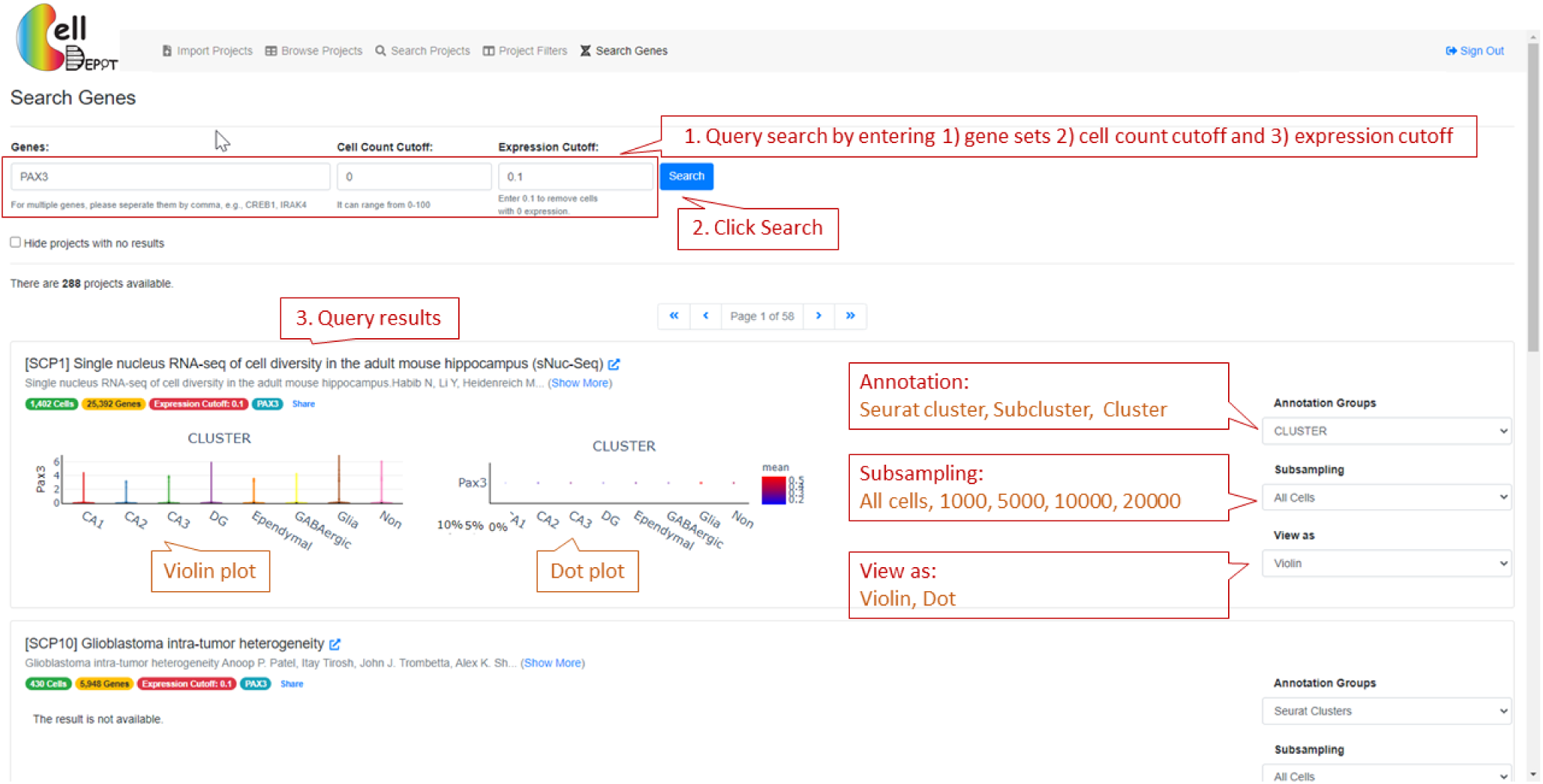
The layout of ‘Search Genens’

#### 6. How to set up cron job?

The following cron job entry is needed to convert h5ad file to CSC format on the background, *@hourly <user-name> cd /var/www/html/celldepot/app/core; php* .*/api_toCSCh5ad*.*php* Please make sure that the user has the permission to write in the data directory.

